# The extinction crisis of the world’s amphibians is easing rapidly

**DOI:** 10.1101/2023.11.22.568382

**Authors:** Martijn van de Pol

## Abstract

A recent paper (*Nature* 622, 308–14)^1^ analysed changes in the extinction risk of 7,102 amphibian species from 1980 to 2004 to 2022 using the Red List Index^2,3^ (RLI) of the International Union for the Conservation of Nature (IUCN). It concluded that the global amphibian extinction crisis has not abated. However, this conclusion appears too pessimistic and not supported by the data. Rather, it is a result of intrinsic limitations of the RLI. These limitations have implications well beyond amphibians, as the RLI is used worldwide for policy, for example as a headline indicator for the Kunming-Montreal Global Biodiversity Framework^4^.

## Large progress in halting the amphibian extinction crisis

A reduced RLI in the most recent assessment of 2022 indeed shows that the number of threatened amphibian species is still increasing worldwide^1^. However, using a more informative integration of changes in rates of up/downlisting^similar to 5^, I calculated that the underlying extinction rate has decreased threefold over this period: 0.00153/year in 1980-2004 to 0.00052/year in 2004-22 period (Extended Methods). Such a reduction in extinction rate implies that the expected time to extinction has improved from a few centuries to a few millennia over the past decades for endangered amphibians (Fig 1A). The extinction crisis thus appears to be easing rapidly.

**Figure 1.**
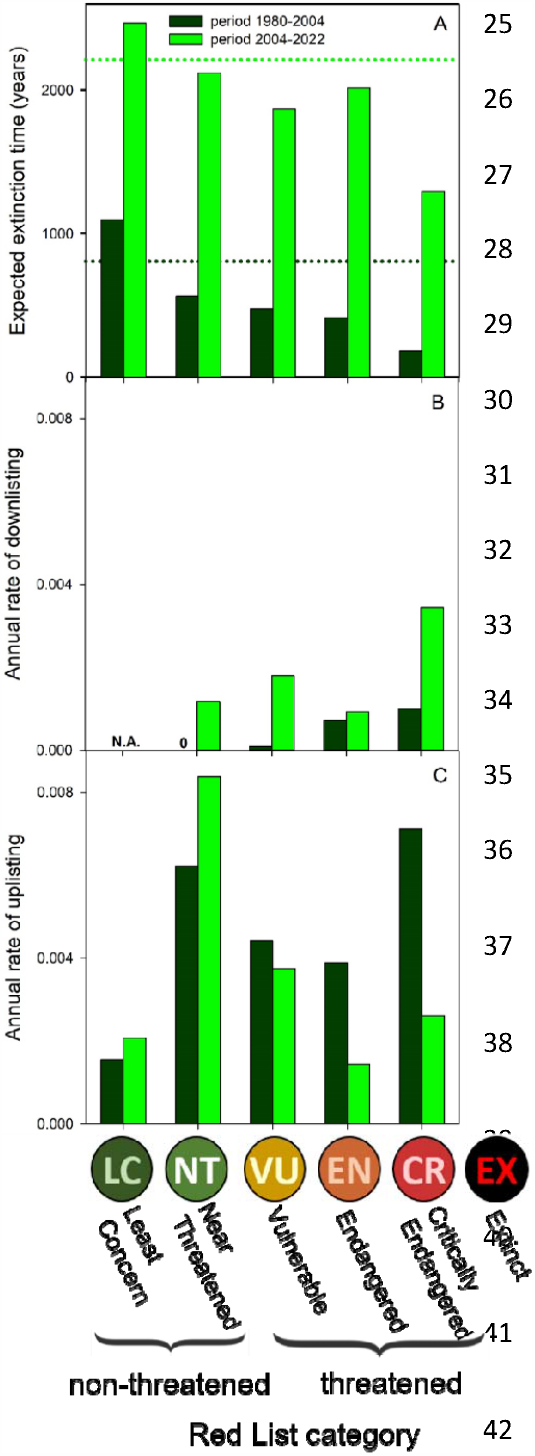
Changes over time in (A) expected extinction time, (B) annual rates of down-, and (C) up-listing of amphibians in specific Red List Categories. Extinction times, downlisting rates and most uplisting rates (but not for LC and NT) greatly improved from the period 1980-2004 to 2004-2022. Downlisting means a species has changed to a less threatened Red List category implying that its extinction risk has decreased (improved). In (A) the horizontal dotted lines show the average extinction time across all amphibian species.

Further analysis of changes in transition rates over time provides more good news. The rates of genuine downlisting overwhelmingly increased over time (Fig. 1B), while the rates of uplisting decreased over time for species in the threatened categories (Fig. 1C). Thus, both the rates at which species are recovering and become threatened are generally improving. The observation that these rates particularly improved for threatened species strongly suggests that conservation efforts played a key role. Indeed, the number of downlistings that were attributed to conservation efforts more than doubled from 0.96/yr in the period 1980-2004 to 2.22/yr in the period 2004-22 (Extended Methods).

My results imply that conservation efforts are increasingly making a difference, another positive development overlooked by Luedtke et al^1^. Overall, this gives reason for some careful optimism even if we have not reached Nature Positive yet (i.e. extinction rates are still well above background rates of one in a million years^6^). Large progress has been made over a short period of only a few decades, and it is important that this is acknowledged because we can build on it for the future to stop the amphibian decline completely.

### Limitations of the influential Red List Index

The RLI is an easily applicable metric that is widely used for comparison among periods, taxonomic groups or geographic regions^7–11^. Consequently, RLI is used worldwide for tracking progress towards halting biodiversity loss and designing–and evaluating the effectiveness of—biodiversity management and policy^4,9,12–15^. However, my analysis highlights that its use can obscure large changes in extinction rates over time. There are at least two issues with the RLI.

First, the RLI weights a change from LC to NT equally heavy as one from NT to VU, or from CR to EX^2^ (Fig. 1 explains abbreviations). This ignores that some changes in threat category increase the extinction risk of a species much more than others. For example, for amphibians a change from LC to NT (or EN to CR) roughly halved a species’ expected time to extinction, while a change from NT to VU (or EN to VU) barely influenced a species’ extinction time (Fig. 1A, period 1980-2004). A similar nonlinear pattern has been reported for birds^5^. The equidistant weights used in RLI thus poorly reflect real-world changes in extinction risk.

Second, differences among periods (or regions, species groups) in RLI are not only caused by rates of extinction and up/downlisting varying among periods, but also by differences in the initial distribution of species among Red List categories. Ultimately, we want to understand what causes underlying rates to vary, because if we know how to improve these, then the Red List status will follow. However, changes or differences in RLI also depend on how many species are already threatened, which is a legacy of extinction rates in previous periods.

This legacy component of RLI can lead to ambiguous results, which becomes obvious from a simple thought experiment: Let’s assume we compare three groups of species over time, from different regions, or taxa^as in Fig. 3 in 1^. All groups have the same fixed annual rates of up and downlisting. However, group 1 consists of only VU species, while group 2 consist of 50% LC and 50% CR, and group 3 consist of 40% LC and 60% CR species. Consequently, groups 1 and 2 have the same RLI at the start (0.6), while group 3 has the lowest RLI of 0.52. We thus expect group 3 to have the highest extinction risk based on their RLI. Furthermore, given that groups have the same rates of up and downlisting and that rates do not change in our experiment, the underlying extinction risk also does not change over time. If RLI is a useful proxy of extinction risk, then the relative differences in RLI among groups should persist.

However, if we follow these groups over time, the RLI of group 1 declines fastest, overtaking group 3 as the group with the lowest RLI (Fig. 2). Based on the RLI we would conclude that threats have increased much more for group 1 than group 3, and that group 1 is now most at risk. However, no risks changed in our experiment for these groups with identical rates. Also, groups 1 and 2 had the same RLI at the start (supposedly reflecting equivalent extinction risk) and both up and downlisted at the same constant rates. Nonetheless, the change in RLI over time decelerates for group 1, while it accelerates for group 2 over time. Interpreting accelerations versus decelerations in RLI over time as evidence for groups having experienced different changes in risk and threats over time is thus fraught with problems.

**Fig. 2:**
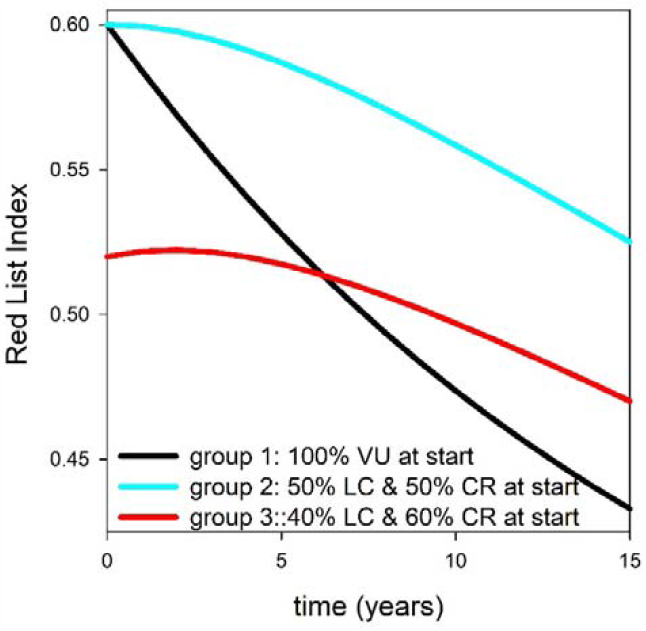
RLI changes differently over time among groups, even when rates of up and downlisting do not change over time and are the same for groups. Groups only differed in their initial distribution of Red List categories (see legend; VU=Vulnerable, LC=Least Concern, CR=Critically Endangered). Rates of up and downlisting were equated to those of amphibians in the period 2004-22 (Extended Methods).

Both limitations of RLI have not only obfuscated that amphibian extinction risk eased rapidly but can also lead to the misidentification of drivers of extinction. For example, according to Luedtke *et al*.^1^ the rate of decline of salamanders and newts accelerated in the period 2004-22, while it decelerated for frogs. They attributed such opposing changes in RLI to a difference in the timing of disease impact for these groups. However, I calculated that the extinction rate of both groups decelerated in the 2004-22 period (Fig. Extended Data), suggesting that disease impact has not increased recently in salamanders and newts at all.

The IUCN advocates using RLI to quantify changes in extinction risk over time and compare this for regions, or species groups relevant to policy mechanisms^12^. However, my study highlights that much caution is needed when using RLI. Analyses of transition rates among threat classes can provide additional—and likely more robust and meaningful—insights^5, this study^. Here, such analysis provided a more optimistic view of the amphibian extinction crisis.

## Extended Methods

### Red List data and categories

Comprehensive IUCN Red Lists were available for the years 1980, 2004 and 2022 for 8,011 amphibian species, see supplement Luedtke et al.^1^. The IUCN Red List considers four categories in addition to those listed in Fig. 1. Species classified as Extinct in the Wild (EW, 0.012% of all amphibian species), as Critically endangered – Probably Extinct in the Wild (CR(PEW), 1.5%) and as Critically endangered – Probably Extinct (CR(PE), 0.004%) were considered equivalent to Extinct (EX) here, similar as done in the RLI calculation^2^. Data Deficient (DD, 11.3%) species are excluded from RLI calculation^2^, and to allow for direct comparison with Luedtke et al.^1^, I also omitted DD species here.

By omitting a substantial number of DD species, the RLI likely underestimates extinction risk. The reason is that DD species are predicted to have a higher extinction risk than the Data Sufficient (DS, i.e. non-DD) species included in the RLI^16,17^. Furthermore, to enable comparisons of RLI over time, DD that became DS at a later point in time were back-casted by Luedtke et al.^1^ for previous Red Lists based on their later DS category and any reported trends in numbers and threats over time^1^. How the assumptions made in back-casting influence the RLI changes over time remains to be explored. The methodology described below based on transition rates can readily include the DD category and would not require back-casting, but I did not consider this here, as it would make the comparison with RLI results hard to interpret.

### Rates of down and uplisting

I calculated the proportion of individuals that moved from a given category to another category in the next Red List. This gives us a transition matrix *P* for the period 1984-2004 and one for the period 2004-2022. As these periods have different lengths, I next transformed each transition matrix into an annual transition matrix *Q*. The transformation to annual probabilities is straightforward once we consider that any square matrix can be diagonalized such that *Q*= *VDV*^-1^, where *V* is the matrix of the right eigenvectors of matrix *Q* and *D* is a diagonal matrix with the eigenvalues of matrix *Q*. Here *Q* is unknown, but we know that P= *Q*^*k*^ = *VD*^*K*^*V*^-1^, where *k* is the number of years between two Red Lists. As the eigenvectors and eigenvalues of *Q* and *Q*^*k*^ are identical we can calculate *V* and *D* from matrix *P*, and use this to calculate *Q*.

### Extinction times and rates

Expected extinction times can be calculated as the inverse of the annual extinction rate across all species considered^5^. Annual extinction rates can be calculated in various ways from the annual transition matrix *Q*. A previous study^5^ calculated the average extinction rate as the inverse of the hitting time of the absorbing category EX of matrix *Q*_*T*_, the transient segment of matrix *Q*. Another option is to calculate the asymptotic (long-term) extinction rate as 1-, *λ* where *λ* is the dominant eigenvalue of matrix *Q*_*T*_^18^. Both methods yielded very similar results in terms of temporal changes in extinction rates (see code and analysis in Supplementary file). In my analysis I focus on the latter method, as the asymptotic extinction rate is independent of the initial distribution of species across all categories, and thus changes over time or among groups of species are not confounded with differences in the initial distribution. It should be noted that the above methods calculate the extinction rate from the entire annual transition matrix, not just from the transitions into the absorbing Extinct category.

### Thought experiment

The three groups in the experiment each had a different initial distribution of species across the Red List categories LC, NT, VU, EN, CR, and EX: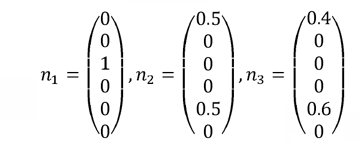. All groups had the same transition matrix across categories *Q* that was constant over time, with values equal to the previously calculated values of *Q* of amphibians over the period 2004-2022. By iterating *n*_*x*_ (*t*+1) = *Q* × *n*_*x*_ (*t*) one obtains the distribution of species across categories over time and from this the RLI at each timestep can be calculated.

## Supporting information

PDF of R code, analysis and all results

## Data and Code Availability

All data is available from the Supplement of Luedtke et al.^1^. My code (using R version 4.2.3)^19^ used to reanalyse their data and produce all results is available as a Supplementary File.

### The author declares that no competing interests exist

## Extended Data Figure

**Extended Data Figure:**
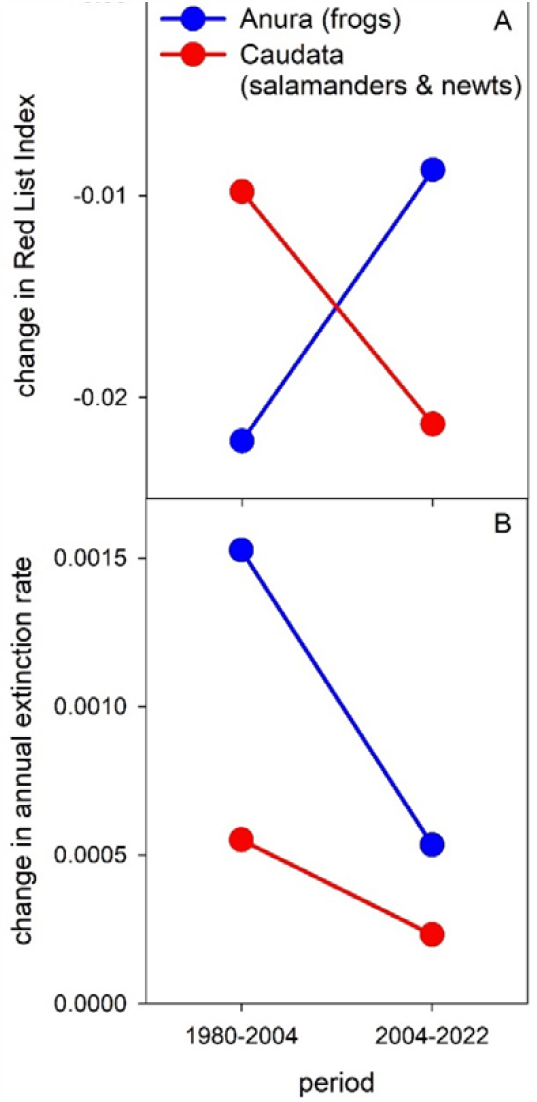
Contrasting changes in threat risk over time for Anura (frogs) and Caudata (salamanders & newts) according to (A) the Red List Index and (B) annual the rate of extinction. (A) is adapted from Fig. 3d in. Luedtke et al.^1^

