## Supplementary material for "The extinction crisis of the world’s amphibians is easing rapidly": PDF of R code, analysis and all results

### Analysis belonging to 'The extinction crisis of the world's amphibians is easing rapidly', a re-analysis of the paper 'Ongoing declines for the world's amphibians in the face of emerging threats' by Leudtke et al. Nature 622, 308–14

Martijn van de Pol - James Cook University

2023-11-18

#### R Markdown

This is an R Markdown document. Markdown is a simple formatting syntax for authoring HTML, PDF, and MS Word documents. For more details on using R Markdown see http://rmarkdown.rstudio.com.

#### Goal

in this R document I re-analyse the changes in threat status over time (lists 1980, 2004 and 2022) from the paper 'Ongoing declines for the world's amphibians in the face of emerging threats (Nature 622, 308–14)' to show that:

1. extinction rate has decreased threefold over this period and expected extinction times similarly
2. most rates of uplisting and downlisting have improved over time, and that conservation effort played a key role in this improvement
3. an eperiment to illsustrate that changes in the RLI voer time are hard to interpret as they depends on legacy effects
4. that the extinction risk of salamanders and newtshas not accelerated over time (the reduction in RLI has accelerated, but not their extinction rate).

```
library(openxlsx)
# below file is from the supplement of the original paper 'Ongoing declines for the world's amphibians
amphib<-read.xlsx(xlsxFile = "All_amphibians_tabular_data.xlsx", sheet = 1, skipEmptyRows = FALSE)
# rename the column names where key data is stored
amphib$'1980'<-amphib$'1980.GAA2.Red.List.Category'
amphib$'2004'<-amphib$'2004.GAA2.Red.List.Category'
amphib$'2022'<-amphib$'2022.GAA2.Red.List.Category'
```

#### Functions

Below are five functions that:

1. Create a transiton matrix between all states derived from two consecutive red lists (1980-2004 or from 2004-2022
2. Recalculate the transition matrix over a period between two IUCN list to annual transition probabilities using standard matrix algebra

3. Calculate the asymptotic (long-term) extinction rate of the transition matrix using matrix algebra, code from Vindenes, Y., Le Coeur, C., & Caswell, H. (2021). Introduction to matrix population models. Demographic Methods across the Tree of Life. Oxford University Press, New York, 163-181.
4. Calculate the expected extinction rate and time to extinction using matrix algebra and Markov theory (code translated from MatLab code from Monroe MJ, Butchart SHM, Mooers AO, Bokma F. 2019 The dynamics underlying avian extinction trajectories forecast a wave of extinctions. Biol. Lett. 15: 20190633. http://dx.doi.org/10.1098/rsbl.2019.0633)) whom previously used this method on avian Red List data.
5. Calculate the Red List Index (as described in Butchart SHM, Akc¸akaya HR, Chanson J, Baillie JEM, Collen B, et al (2007) Improvements to the Red List Index. PLoS ONE 2(1): e140. doi: 10.1371/journal.pone.0000140)

**function 1: create a transition matrix between threat states of two subequent IUCN lists**

```
construct_matrix<-function(data, start, end) {
  states<-c("DD", "LC", "NT", "VU", "EN", "CR",  "EX")
  # recode some specific categories into above states
  data[which(data[,which(colnames(data)==start)]=="EW") ,which(colnames(data)==start)]<-"EX"
  data[which(data[,which(colnames(data)==end)]=="EW") ,which(colnames(data)==end)]<-"EX"
  data[which(data[,which(colnames(data)==start)]=="CR(PE)") ,which(colnames(data)==start)]<-"EX"
  data[which(data[,which(colnames(data)==end)]=="CR(PE)") ,which(colnames(data)==end)]<-"EX"
  data[which(data[,which(colnames(data)==start)]=="CR(PEW)") ,which(colnames(data)==start)]<-"EX"
  data[which(data[,which(colnames(data)==end)]=="CR(PEW)") ,which(colnames(data)==end)]<-"EX"
  # define matrix
  mat<-matrix(data=0, nrow=length(states), ncol=length(states), dimnames = list(states,states))
  n0<-matrix(data=0, nrow=1, ncol=length(states), dimnames = list("N",states))
  for(i in 1:length(data[ ,which(colnames(data)==start)])) {
    mat[data[i, which(colnames(data)==end)], data[i ,which(colnames(data)==start)]]<-(1+ mat[data[i,whic
    n0[1,data[i ,which(colnames(data)==start)]]<-(1+n0[1,data[i ,which(colnames(data)==start)]])
  }
  # calculate transition matrix
  for (i in 1:dim(mat)[1]) {
    mat[,i]<-mat[,i]/sum(mat[,i])
  }
 return(list(mat, n0))
}
```

**function 2: recalculate the transition matrix over a period between two IUCN list to annual transition probabilities using standard matrix algebra**

```
annual_transition<-function(A, duration) {
  # we have a transition matrix A over a certain period, say k years between two IUCN red list,
  # we now want to know what the annual transition matrix is, such that we can compare it to other peri
  # this issue is relevant for amphibians as list 1 and 2 were 24 years apart (2004-1980) while list 2
  # to calculate the annual transition matrix from a trasnsiton matrix spanning a longer period we use
  # 1. We first diagonalize matrix Q: Q=V%*%D%*%V^-1, where D is the diagonal matrix of Q with the eige
  # 2. if we take Q*Q=VDV^-1VDV^-1=VD^2V^-1, which means that Q^k=V%*%D^k%*%V^-1, where k is the number
  # 3. the V (right eigenvectors) of Q and of Q^k are identical, so V of Q can be calculated from Q^k,
  # 4. The diagonal matrix D^k can be calculated from Q^k=A, and since k is also known this means we ca
  # 5. This means that Q can be reconstructed using Q=V%*%D%*%V^-1
  ev_power <- eigen(A)
  P_power<-ev_power$vectors
```

```
  V_power<-diag(ev_power$values^(1/duration))
  P_powerinv<-solve(P_power)
  A1<-P_power%*%V_power%*%P_powerinv
  A1[which(Re(A1)<0)]<-0  # this is rounding very small negative numbers (due to rounding errors) to ze
  for(i in 1:(dim(A)[1])) { A1[,i]<-A1[,i]/sum(A1[,i]) } # this is ensuring that all transitions sum up
  colnames(A1)<-colnames(A)
  rownames(A1)<-rownames(A)
  return(A1)
}
```

**function 3: calculate the asymptotic extinction rate of a transition matrix (with extinction rate equaling 1-growth rate lambda)**

```
 wvlambda <- function(MatA){
 ev <- eigen(MatA)
 tev <- eigen(t(MatA))
 lmax<-which.max(Re(ev$values))
 W <- ev$vectors
 V <- tev$vectors
 w <- as.matrix(abs(Re(W[, lmax]))/sum(abs(Re(W[, lmax]))))
 v <- as.matrix(abs(Re(V[, lmax])))
 v <- v/sum(v*w)
 return(list("lambda"=max(Re(ev$values)),"w"=w,"v"=v))
 }
```

**function 4: calculate the expected extinction rate and extinction time**

```
extinct_risk<-function(mat, n0) {
  states<-c("DD", "LC", "NT", "VU", "EN", "CR",  "EX")
  Q<-t(mat)
  R<-Q[-which(colnames(mat)=="DD"),-which(colnames(mat)=="DD")]
  R<-R[-which(colnames(R)=="EX"),-which(colnames(R)=="EX")]
  I<-diag(dim(R)[1])
  temp<-I-R
  F<-solve(temp)   # this takes the inverse
  C<-rep(1,dim(R)[1])
  Tt<-F%*%C  # expected times till extinction for each class (in timestep units of the matrix period)
  n0<-n0[-which(states=="DD" |  states=="EX")]
  K<-t(n0/sum(n0))  # the starting distribution
  time_to_extinction<-K%*%Tt  # average time to extinction
  extinction_risk<-1/time_to_extinction # extinction intensity per timestep (period between two IUCN li
  return(list(extinction_risk,Tt,K))
}
```

**function 5: calculate the Red List Index (RLI)**

```
RLI<-function(data) {
  indexstates<-c("EX","CR","EN","VU", "NT", "LC")
  weights<-c(5,4,3,2,1,0)  # corresponding weights of states
  N<-length(data)-length(which(data=="DD")) # all species excluding DD (and EX before the first year as
  M<-max(weights)*N
  TT<-length(which(data=="LC"))*weights[which(indexstates=="LC")]+
```

```
      length(which(data=="NT"))*weights[which(indexstates=="NT")]+
      length(which(data=="VU"))*weights[which(indexstates=="VU")]+
      length(which(data=="EN"))*weights[which(indexstates=="EN")]+
      length(which(data=="CR"))*weights[which(indexstates=="CR")]+
      (length(which(data=="EX")) +
       length(which(data=="EW")) +
       length(which(data=="CR(PE)"))+
       length(which(data=="CR(PEW)")))*weights[which(indexstates=="EX")] # all species excluding DD, no
  index<-1-(TT/M)
  return(index)
}
```

##### Results 1 : Comparing the exinction rate for the two different period (1980-2004 vs. 2004-2022). Has the extinction rate accelerated, stabilized or slowed down over time? This analysis is for all amphibians combined.

```
# period 1: 1980-2004
start<-1980
end<-2004
mat1<-construct_matrix(amphib, start=start, end=end)[[1]] # construct transition matrixover whole perio
annual_mat1<-annual_transition(A=mat1, duration=(end-start)) # construct annual transition matrix
n01<-construct_matrix(data=amphib, start=start, end=end)[[2]] # calculate the distribution of species
smat1<-annual_mat1[-which(colnames(annual_mat1)=="DD"),-which(colnames(annual_mat1)=="DD")]
smat1<-smat1[-which(colnames(smat1)=="EX"),-which(colnames(smat1)=="EX")]

# period 2: 2004-2022
start<-2004
end<-2022
mat2<-construct_matrix(amphib, start=start, end=end)[[1]] # construct transition matrix over whole peri
annual_mat2<-annual_transition(A=mat2, duration=(end-start)) # construct annual transition matrix
n02<-construct_matrix(data=amphib, start=start, end=end)[[2]] # calculate the distribution of species a
smat2<-annual_mat2[-which(colnames(annual_mat2)=="DD"),-which(colnames(annual_mat2)=="DD")]
smat2<-smat2[-which(colnames(smat2)=="EX"),-which(colnames(smat2)=="EX")]

# calculate asymptotic extinction rates
asympt_exinction_rate1<-1-wvlambda(smat1)$lambda # percentage decline per year over period 1
asympt_exinction_rate2<-1-wvlambda(smat2)$lambda # percentage decline per year over period 1
ratio<-asympt_exinction_rate1/asympt_exinction_rate2
print(c(asympt_exinction_rate1, asympt_exinction_rate2, ratio))
```

```
## [1] 0.0015288254 0.0005236535 2.9195363806
```

We see that in period 1 (1980-2004) the extinction rate was 0.00153/year and in period 2 (2004-2022) it was 0.00052/year, this amounts to a threefold (2.92) decrease in extinction rate over time.

Above method focuses on the asymptotic extinction rate, but we can also calculate the expected extinction rate using methods previously applied to IUCN red list data (Monroe MJ, Butchart SHM, Mooers AO, Bokma F. 2019 The dynamics underlying avian extinction trajectories forecast a wave of extinctions. Biol. Lett. 15: 20190633) to show that my main result is robust to the specific metric of extinction rate used. I also calculate the expected extinction times for both periods for a species in a given Red List category (Fig. 1a in my paper):

```
# period 1: 1980-2004
ex_rate1<-extinct_risk(mat=annual_mat1, n0=n01)[[1]]
```

```r
ex_times1<-extinct_risk(mat=annual_mat1, n0=n01)[[2]]
colnames(ex_times1)<-("expected time to exinction 1980-2004 (years)")
ex_times1_all<-sum(extinct_risk(mat=annual_mat1, n0=n01)[[3]]*t(ex_times1)) # expected extincton time a

# period 2: 2004-2022
ex_rate2<-extinct_risk(mat=annual_mat2, n0=n02)[[1]]
ex_times2<-extinct_risk(mat=annual_mat2, n0=n02)[[2]]
colnames(ex_times2)<-("expected time to exinction 2004-2022 (years)")
ex_times2_all<-sum(extinct_risk(mat=annual_mat2, n0=n01)[[3]]*t(ex_times2)) # expected extincton time a

print(c("annual extinction rate 1980-2004", "annual extinction rate 2004-2022", "ratio of extinction ra
```

```
## [1] "annual extinction rate 1980-2004"
## [2] "annual extinction rate 2004-2022"
## [3] "ratio of extinction rate between periods"
```

```r
print(c(ex_rate1, ex_rate2, ex_rate1/ex_rate2))
```

```
## [1] 0.001248804 0.000454945 2.744955681
```

```r
#compare the expected extinction times across the period for a species in a given state
print(ex_times1)
```

```
##    expected time to exinction 1980-2004 (years)
## LC                                    1098.3520
## NT                                     562.8148
## VU                                     477.5072
## EN                                     416.3597
## CR                                     184.6469
```

```r
print(ex_times2)
```

```
##    expected time to exinction 2004-2022 (years)
## LC                                     2470.350
## NT                                     2121.961
## VU                                     1874.227
## EN                                     2017.829
## CR                                     1296.378
```

```r
#compare the expected extinction times across the period for all amphibian species see Fig. 1a
print(c(ex_times1_all, ex_times2_all))
```

```
## [1]  800.7662 2207.0033
```

We can see that also in this analysis there is a rougly threefold (2.74) reduction in expected extinction rate between the periods 1980-2004 and 2004-2022 (0.0012/year in 1980-2004 vs. 0.00045/year in period 2004-2022). Note that this latter method is possibly less informative as the extinction rate is also affected by the starting distribution across states, while this is not the case for the asymptotic extinction rate above.

The expected extinction times also improved dramatically in period 2004-2022 compared to 1980-2004 period (these values were used to make Fig. 1a). For example for endagngerd species (EN and CR) it changed from 184-416 years (a few centuries) to 1296-2017 years (a few millenia).

#### Result 2: changes in rates of up and downlisting between the two periods (1980-2004 vs. 2004-2022).

```
mat_difference<-annual_mat2-annual_mat1
mat_difference<-mat_difference[-which(colnames(mat_difference)=="DD"),-which(colnames(mat_difference)==
mat_difference<-mat_difference[,-which(colnames(mat_difference)=="EX")]
mat_difference
```

```
##               LC            NT            VU            EN            CR
## LC -5.203711e-04  1.179536e-03  7.785965e-05  8.832903e-05  0.0000000000
## NT  3.612869e-04 -3.330161e-03  1.629968e-03 -1.604615e-04  0.0002774885
## VU  4.926568e-04  6.050898e-05 -1.041908e-03  2.817378e-04  0.0008684667
## EN -1.772087e-04  3.041504e-03 -4.527165e-04  2.242014e-03  0.0012976521
## CR -9.054366e-05 -9.513882e-04  2.600001e-04 -1.384282e-03  0.0020668023
## EX -6.582025e-05  0.000000e+00 -4.732032e-04 -1.067337e-03 -0.0045104096
```

```
uplist_rates<-mat_difference[lower.tri(mat_difference)]
downlist_rates<-mat_difference[upper.tri(mat_difference)]
sign(uplist_rates)
```

```
##  [1]  1  1 -1 -1 -1  1  1 -1  0 -1  1 -1 -1 -1 -1
```

```
sign(downlist_rates)
```

```
##  [1]  1  1  1  1 -1  1  0  1  1  1
```

```
# we can sum these rates per category
states<-c("LC", "NT", "VU", "EN", "CR",  "EX")
updownrates<-data.frame(category = states[-6], uplist_rate1 = rep(NA,length(states[-6])), downlist_rate
#period 1
m1<-annual_mat1[-which(colnames(annual_mat1)=="DD"), -which(colnames(annual_mat1)=="DD")]
updownrates[which(states=="LC"),"uplist_rate1"]<-sum(m1[2:6,which(states=="LC")])
updownrates[which(states=="NT"),"uplist_rate1"]<-sum(m1[3:6,which(states=="NT")])
updownrates[which(states=="VU"),"uplist_rate1"]<-sum(m1[4:6,which(states=="VU")])
updownrates[which(states=="EN"),"uplist_rate1"]<-sum(m1[5:6,which(states=="EN")])
updownrates[which(states=="CR"),"uplist_rate1"]<-sum(m1[6:6,which(states=="CR")])
updownrates[which(states=="LC"),"downlist_rate1"]<-NA
updownrates[which(states=="NT"),"downlist_rate1"]<-sum(m1[1:1,which(states=="NT")])
updownrates[which(states=="VU"),"downlist_rate1"]<-sum(m1[1:2,which(states=="VU")])
updownrates[which(states=="EN"),"downlist_rate1"]<-sum(m1[1:3,which(states=="EN")])
updownrates[which(states=="CR"),"downlist_rate1"]<-sum(m1[1:4,which(states=="CR")])
#period 2
m2<-annual_mat2[-which(colnames(annual_mat1)=="DD"), -which(colnames(annual_mat1)=="DD")]
updownrates[which(states=="LC"),"uplist_rate2"]<-sum(m2[2:6,which(states=="LC")])
updownrates[which(states=="NT"),"uplist_rate2"]<-sum(m2[3:6,which(states=="NT")])
updownrates[which(states=="VU"),"uplist_rate2"]<-sum(m2[4:6,which(states=="VU")])
updownrates[which(states=="EN"),"uplist_rate2"]<-sum(m2[5:6,which(states=="EN")])
updownrates[which(states=="CR"),"uplist_rate2"]<-sum(m2[6:6,which(states=="CR")])
updownrates[which(states=="LC"),"downlist_rate2"]<-NA
updownrates[which(states=="NT"),"downlist_rate2"]<-sum(m2[1:1,which(states=="NT")])
updownrates[which(states=="VU"),"downlist_rate2"]<-sum(m2[1:2,which(states=="VU")])
updownrates[which(states=="EN"),"downlist_rate2"]<-sum(m2[1:3,which(states=="EN")])
updownrates[which(states=="CR"),"downlist_rate2"]<-sum(m2[1:4,which(states=="CR")])
updownrates
```

```
##   category uplist_rate1 downlist_rate1 uplist_rate2 downlist_rate2
```

```
## 1       LC  0.001553591            NA  0.002073962            NA
## 2       NT  0.006222333  0.000000e+00  0.008372957  0.0011795362
## 3       VU  0.004416388  9.122607e-05  0.003750469  0.0017990540
## 4       EN  0.003891546  7.188809e-04  0.001439927  0.0009284862
## 5       CR  0.007122794  9.998762e-04  0.002612384  0.0034434835
```

The matrix mat_difference shows the difference between the two periods in the annual transition probabilities (period 2-period 1). The probabilites of uplisting are in the matrix elements below the diagonal and are improving (negative differences) over time for 9 out of 15 transitions 1 no change, 4 deteriorations). The rates of downlisting are in the matrix elements above the diagonal and are improving (positive differences) for 8 out of 10 transitions (1 did not change, 1 deteriorated). Overall this means that 68% of the transition rates improved over time (17/25) and only 20% deteriorted (5/25).

If we sum(marize) them per category we see that all probabilties are improving from period 1980-2004 to period 2004-2022, except for the rates of uplisting for LC and NT (values from printed table used in Fig. 1b and 1c).

#### How much of the downlisting are due to conservation efforts in both periods?

The Red List has attributed which donwlistings were attributed to have happened due to conservation efforts. Luedtke et al. already mentioned that about half of the donwlistings over both periods could be attributed to conservation efforts. However, they did not analyze whether there were more downlistings due to conservation efforts in the more recent period. I thus analyzed how many of such conservation-driven downlisting occured in both periods.

```
cons_donwlist1<-length(which(amphib$`1980-2004.Genuine.downlisting.due.to.conservation`=="Yes"))/(2004-
cons_donwlist2<-length(which(amphib$`2004-2022.Genuine.downlisting.due.to.conservation`=="Yes"))/(2022-
print(c(cons_donwlist1, cons_donwlist2, cons_donwlist2/cons_donwlist1))
```

```
## [1] 0.9583333 2.2222222 2.3188406
```

We see that there were more than twice (2.32 times) as many donwlistings per year in the more recent period 2004-2022 (2.22 species/year) than in the period 1980-2004 (0.96 species/year).

### Results 3: Thought experiment to show dependency of RLI on the initial distribution of species

```
# this experiment considered three groups with varying starting distribution but with the same constant
# we take starting distribution of larval developers, states in below vectors are respectively: LC, NT,
n1_exp<-c(0,0,1,0,0,0)  # all species are Vulnerable
n2_exp<-c(0.5,0,0,0,0.5,0) # 50% species are Least Concern and 50% Critically Endangered
n3_exp<-c(0.4,0,0,0,0.6,0) # 40% species are Least Concern and 60% Critically Endangered
RLI_1<-RLI_2<-RLI_3<-c()
RLI_1[1]<-sum(n1_exp*rev(seq(from=0,to=1,by=.2)))/sum(n1_exp) # this is a simpler but equivalent way of
RLI_2[1]<-sum(n2_exp*rev(seq(from=0,to=1,by=.2)))/sum(n2_exp)
RLI_3[1]<-sum(n3_exp*rev(seq(from=0,to=1,by=.2)))/sum(n3_exp)

# for the transition matrix we take the values of amphibians in the most recent period 2004-2022 (this
Q<-mat2[-which(colnames(mat2)=="DD"),-which(colnames(mat2)=="DD")]

timesteps<-20
n1_exp_t<-n2_exp_t<-n3_exp_t<-matrix(data=NA, nrow=(timesteps+1), ncol=6, dimnames=list( seq(1,(timestep
n1_exp_t[1,]<-n1_exp
```

```
n2_exp_t[1,]<-n2_exp
n3_exp_t[1,]<-n3_exp

for(i in 1:timesteps) {
    n1_exp_t[i+1,]<-Q%*%n1_exp_t[i,]
    n2_exp_t[i+1,]<-Q%*%n2_exp_t[i,]
    n3_exp_t[i+1,]<-Q%*%n3_exp_t[i,]
    RLI_1[i+1]<-sum(n1_exp_t[i+1,]*rev(seq(from=0,to=1,by=.2)))/sum(n1_exp_t[i+1,]) # this is a simpler
    RLI_2[i+1]<-sum(n2_exp_t[i+1,]*rev(seq(from=0,to=1,by=.2)))/sum(n2_exp_t[i+1,])
    RLI_3[i+1]<-sum(n3_exp_t[i+1,]*rev(seq(from=0,to=1,by=.2)))/sum(n3_exp_t[i+1,])
}

plot(RLI_3, xlim=c(0,15), ylim=c(0.43,0.6), type="l", col="red")
lines(RLI_2, col="blue")
lines(RLI_1, col="black")
```

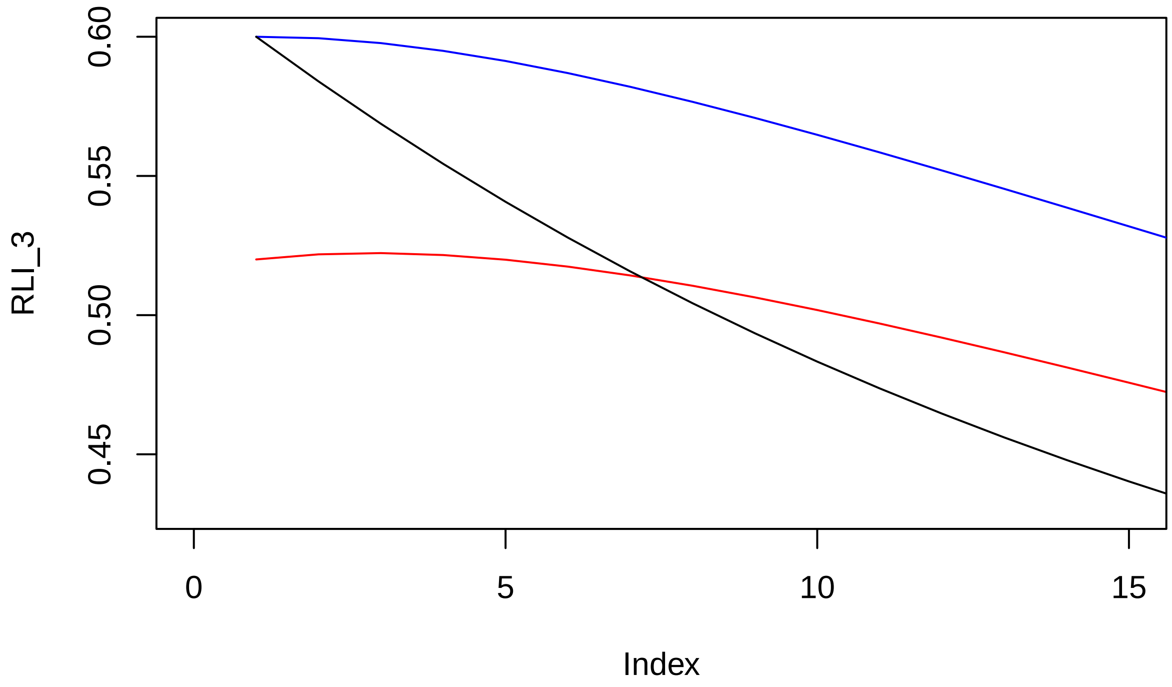


This plot reproduces Fig. 2 in my paper. Groups 1 (black) and 2 (blue) have the same RLI at the start (0.6), while group 3 (red) has the lowest RLI of 0.52. We thus expect group 3 to have the highest extinction risk based on their RLI. Furthermore, given that groups have the same rates of up and downlisting and that rates do not change in our experiment, the underlying extinction risk also does not change over time. If RLI is a useful proxy of extinction risk, then the relative differences in RLI among groups should persist. However, if we follow these groups over time, the RLI of group 1 declines fastest, overtaking group 3 as the group with the lowest RLI (plot above). Based on the RLI we would conclude that threats have increased much more for group 1 than group 3, and that group 1 is now most at risk. However, no risks changed in our experiment for these groups with identical rates. Also, groups 1 and 2 had the same RLI at the start (supposedly reflecting equivalent extinction risk) and both up and downlisted at the same constant rates. Nonetheless, the change in RLI over time decelerates for group 1, while it accelerates for group 2 over time. Interpreting accelerations versus decelerations in RLI over time as evidence for groups having experienced different changes in risk and threats over time is thus fraught with problems.

#### Results 4: Do Caudata (salamanders and newts) and Anura (frogs) differ in their changes in extinction risk over time?

```
amphib_anura<-amphib[which(amphib$Order=="ANURA"),]
amphib_caudata<-amphib[which(amphib$Order=="CAUDATA"),]

#The following results were used to produce the Extended Data Fig.
# Extended Data Fig. Fig.A reproduces the results of Fig 3d in Luedtke et al., but with two differences
# (a) I  ignored the small group of Gymnophiona as the Luedtke et al. also focusses on Caudata and Anur
# (b) instead of plotting RLI for the three points in time (1980-2004-2022) and look at how the RLI cha
delta_anura_period1<-RLI(amphib_anura$'2004')-RLI(amphib_anura$'1980')
delta_anura_period2<-RLI(amphib_anura$'2022')-RLI(amphib_anura$'2004')
delta_caudata_period1<-RLI(amphib_caudata$'2004')-RLI(amphib_caudata$'1980')
delta_caudata_period2<-RLI(amphib_caudata$'2022')-RLI(amphib_caudata$'2004')

print("change in RLI in both periods for anura")
```

```
## [1] "change in RLI in both periods for anura"
```

```
print(c(delta_anura_period1,delta_anura_period2))
```

```
## [1] -0.02215493 -0.00873446
```

```
print("change in RLI in both periods for caudata")
```

```
## [1] "change in RLI in both periods for caudata"
```

```
print(c(delta_caudata_period1,delta_caudata_period2))
```

```
## [1] -0.009817672 -0.021318373
```

```
# Next we calculated the change in extinction rate between the two period to construct Fig. 2b

# period 1: 1980-2004
start<-1980
end<-2004
# anura species
mat1_l<-construct_matrix(amphib_anura, start=start, end=end)[[1]] # construct transition matrix
annual_mat1_l<-annual_transition(A=mat1_l, duration=(end-start)) # construct annual transition matrix
n01_l<-construct_matrix(data=amphib_anura, start=start, end=end)[[2]] # calculate the distribution of s
smat1_l<-annual_mat1_l[-which(colnames(annual_mat1_l)=="DD"),-which(colnames(annual_mat1_l)=="DD")]
smat1_l<-smat1_l[-which(colnames(smat1_l)=="EX"),-which(colnames(smat1_l)=="EX")]
l1_l<-1-wvlambda(smat1_l)$lambda # percentage decline per year over period 1
ex_rate1_l<-extinct_risk(mat=mat1_l, n0=n01_l)[[1]]/(end-start)

# caudata species
mat1_n<-construct_matrix(amphib_caudata, start=start, end=end)[[1]] # construct transition matrix
annual_mat1_n<-annual_transition(A=mat1_n, duration=(end-start)) # construct annual transition matrix
n01_n<-construct_matrix(data=amphib_caudata, start=start, end=end)[[2]] # calculate the distribution of
smat1_n<-annual_mat1_n[-which(colnames(annual_mat1_n)=="DD"),-which(colnames(annual_mat1_n)=="DD")]
smat1_n<-smat1_n[-which(colnames(smat1_n)=="EX"),-which(colnames(smat1_n)=="EX")]
```

```r
l1_n<-1-wvlambda(smat1_n)$lambda # percentage decline per year over period 1
ex_rate1_n<-extinct_risk(mat=mat1_n, n0=n01_n)[[1]]/(end-start)

# period 2: 1980-2004
start<-2004
end<-2022
# anura species
mat2_l<-construct_matrix(amphib_anura, start=start, end=end)[[1]] # construct transition matrix
annual_mat2_l<-annual_transition(A=mat2_l, duration=(end-start)) # construct annual transition matrix
n02_l<-construct_matrix(data=amphib_anura, start=start, end=end)[[2]] # calculate the distribution of sp
smat2_l<-annual_mat2_l[-which(colnames(annual_mat2_l)=="DD"),-which(colnames(annual_mat2_l)=="DD")]
smat2_l<-smat2_l[-which(colnames(smat2_l)=="EX"),-which(colnames(smat2_l)=="EX")]
l2_l<-1-wvlambda(smat2_l)$lambda # percentage decline per year over period 1
ex_rate2_l<-extinct_risk(mat=mat2_l, n0=n02_l)[[1]]/(end-start)

# caudata species
mat2_n<-construct_matrix(amphib_caudata, start=start, end=end)[[1]] # construct transition matrix
annual_mat2_n<-annual_transition(A=mat2_n, duration=(end-start)) # construct annual transition matrix
n02_n<-construct_matrix(data=amphib_caudata, start=start, end=end)[[2]] # calculate the distribution of
smat2_n<-annual_mat2_n[-which(colnames(annual_mat2_n)=="DD"),-which(colnames(annual_mat2_n)=="DD")]
smat2_n<-smat2_n[-which(colnames(smat2_n)=="EX"),-which(colnames(smat2_n)=="EX")]
l2_n<-1-wvlambda(smat2_n)$lambda # percentage decline per year over period 1
ex_rate2_n<-extinct_risk(mat=mat2_n, n0=n02_n)[[1]]/(end-start)

print("asymptotic extintion rate anura period 1, period 2 and difference")
```

```
## [1] "asymptotic extintion rate anura period 1, period 2 and difference"
```

```r
print(c(l1_l, l2_l, l2_l-l1_l))
```

```
## [1]  0.0015265340  0.0005341256 -0.0009924083
```

```r
print("asymptotic extintion rate caudata period 1, period 2 and difference")
```

```
## [1] "asymptotic extintion rate caudata period 1, period 2 and difference"
```

```r
print(c(l1_n, l2_n, l2_n-l1_n))
```

```
## [1]  0.0005510885  0.0002319196 -0.0003191690
```

```r
# this result is robust to the way extinction rate is calculated, if we would use the method of Monroe
print("expected extintion rate anura period 1, period 2 and difference")
```

```
## [1] "expected extintion rate anura period 1, period 2 and difference"
```

```r
print(c(ex_rate1_l, ex_rate2_l, ex_rate2_l-ex_rate1_l))
```

```
## [1]  0.0013020910  0.0004392183 -0.0008628727
```

```r
print("expected extintion rate anura period 1, period 2 and difference")
```

```
## [1] "expected extintion rate anura period 1, period 2 and difference"
```

```r
print(c(ex_rate1_n,ex_rate2_n, ex_rate2_n-ex_rate1_n))
```

```
## [1]  0.0004696736  0.0002181964 -0.0002514773
```

As already shown by Ludtke et al. we see that the change in RLI became less negative (slowed down, -0.0221 vs. -0.0087 ) in period 2 for Anura species (frogs) and that the change in RLI became more negative (accelerated -0.0098 vs. -0.0213) in period 2 for Caudata species (salamanders & newts). This is plotted in Extended Data Fig A.

By contrast, the asymptotic rate of extinction slowed down more than twofold for Caudata in period 2 (0.00055 vs. 0.00023), which is in sharp contrast to the acceleration of the reduction in RLI for Caudata. This slowing down in extinction rate is plotted in Extended Data Fig. B together with the slowing down of extinction rate for Anura in both periods.
